## Supplemental Figures and Tables for "Hidden paths to endless forms most wonderful: Ecology latently shapes evolution of multicellular development in predatory bacteria"

### SUPPLEMENTARY INFORMATION

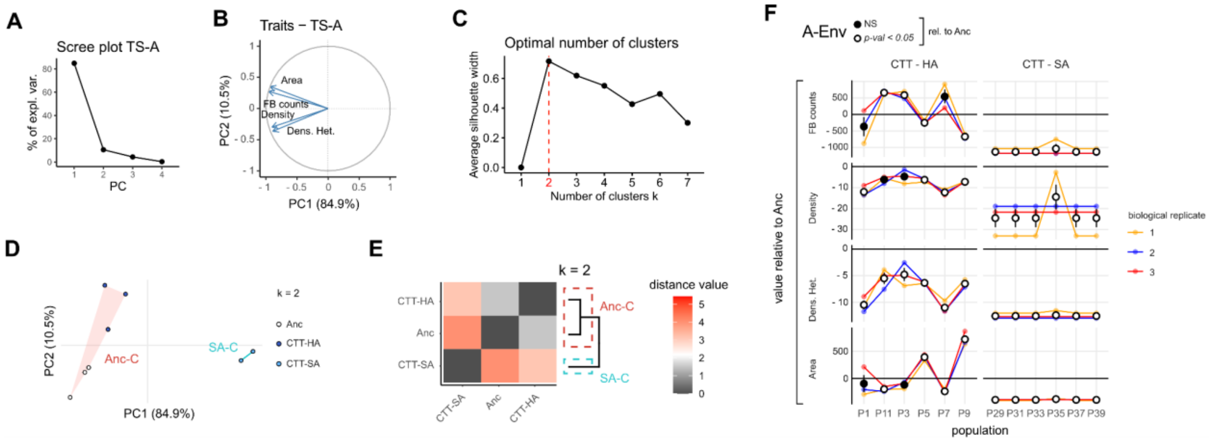


**Fig. S1. PCA, k-means, phenotype-dendrogram and population-level single traits analyses of TS-A.**

**A**, Scree plot indicating the percentage of the explained variance by each principal component obtained from the PCA run on TS-A and Anc together. **B**, Distribution in multivariate space of eigenvectors representing the four morphological traits used in the PCA. **C**, Optimal number of clusters (k = 2) obtained from the average silhouette width (see Methods) of PCA results. **D**, Subdivision of all individual replicates (Anc: open circles, CTT-HA: dark blue circles, CTT-SA: light blue circles) by the two clusters Anc-C (Ancestor-Cluster, red-shaded region) and SA-C (Soft Agar-Cluster, blue-shaded region) obtained from the k-means cluster analysis (k = 2, see Methods). **E**, Heatmap of pairwise Euclidean distances measured between individual centroids mapped on the multivariate space (Fig. 1B). Individual tails color ranging from 0 (grey) to 5 (red) indicate distance values. Dendrogram of hierarchical clustering (see Methods) across all pairwise Euclidean distances is reported on the right side. To facilitate the contrast between the two clustering methods, k-means clusters Anc-C (red-dashed box) and SA-C (light-blue dashed box) (see **D**) were superimposed on the dendrogram. **F**, Single developmental trait values relative to Anc (horizontal-black line set at zero) of all single evolved populations from TS-A. Values were averaged across three independent biological replicates (small colored circles with connecting lines, n = 3). Average values (large circles) are reported with the associated SEM in all cases. Colored lines connect different population values measured within the same biological replicate. In both panels, white and black filled circles indicate average values with a significant or non-significant difference from the Anc, respectively. In **F**, significance levels were calculated with one-way ANOVA followed by two-tailed Tukey tests. *p*-values of all comparisons of evolved treatments with Anc, as well as all pairwise comparisons between evolved treatments are reported in Data Fig. S1.

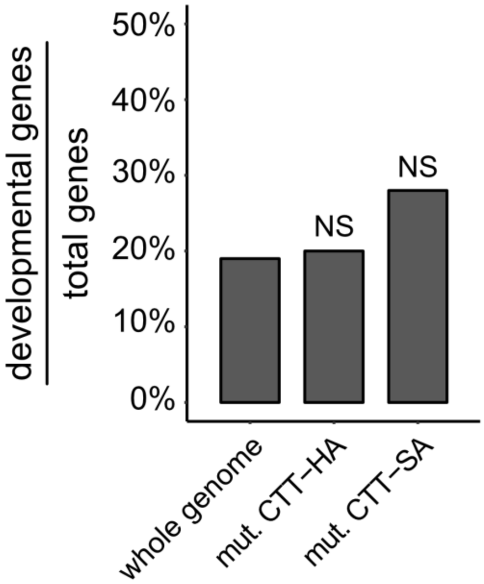


**Fig. S2. Developmental genes are not over-represented among mutated genes in the CTT-HA and CTT-SA treatments.**

Bar graph showing the proportion of all annotated *M. xanthus* genes designated as developmental genes[76] compared to the proportion of developmental genes among genes mutated in the MyxEE-3 cycle 40 CTT-HA and CTT-SA evolved populations[74] (Significance was tested with Fisher's exact test for multiple comparison; NS = not significant). **Data Fig. S2**.

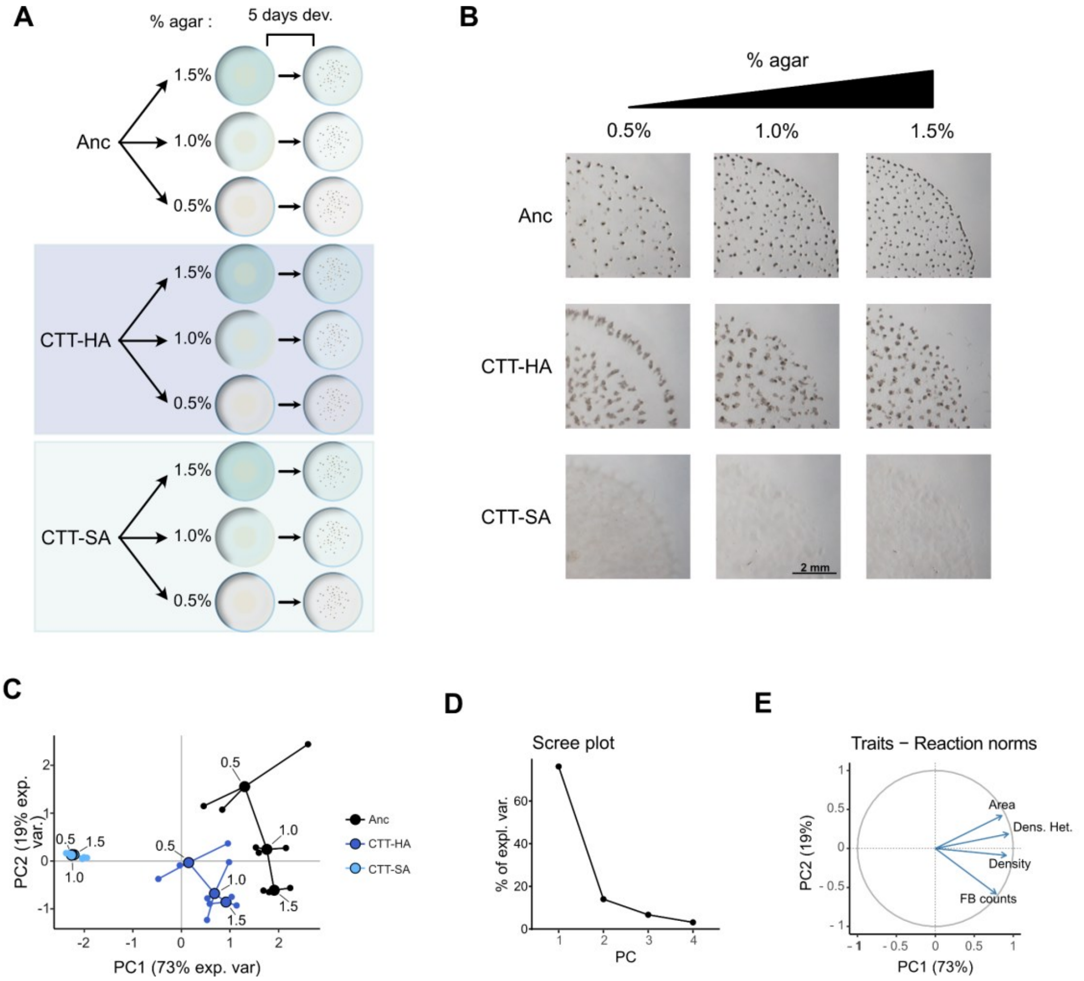


**Fig. S3. Reaction-norm analysis: design, representative images and PCA.**

**A**, Schematic representation of the experimental procedure used to analyze developmental plasticity across the Anc clones and TS-A evolved populations. Following the protocol to induce FB development (see Methods), cells from all evolved populations of both CTT-HA and CTT-SA selective environments and all Anc sub-clones were plated as on TPM starvation plates that differed only in agar concentration along a gradient (0.5%, 1.0% and 1.5%). After five days of starvation, populations were morphologically analyzed. **B**, Images of developmental plates after five days of starvation (CTT-HA = P3; CTT-SA = 39) (scale bar = 2 mm). **C**, PCA of developmental reaction norms across all three agar concentrations. Large circles depict the centroids of three independent replicates (small circles and connecting lines). Anc, CTT-HA and CTT-SA multivariate reaction norms are shown in black, dark blue and light blue, respectively. The x and y axis labels report the percentage of variance explained by the two principal components PC1 and PC2, respectively. **D**, Scree plot indicating variance percentages explained by the four principal components. **E**, Eigenvectors of the four developmental traits used in the PCA. [Data Fig. S3](#).

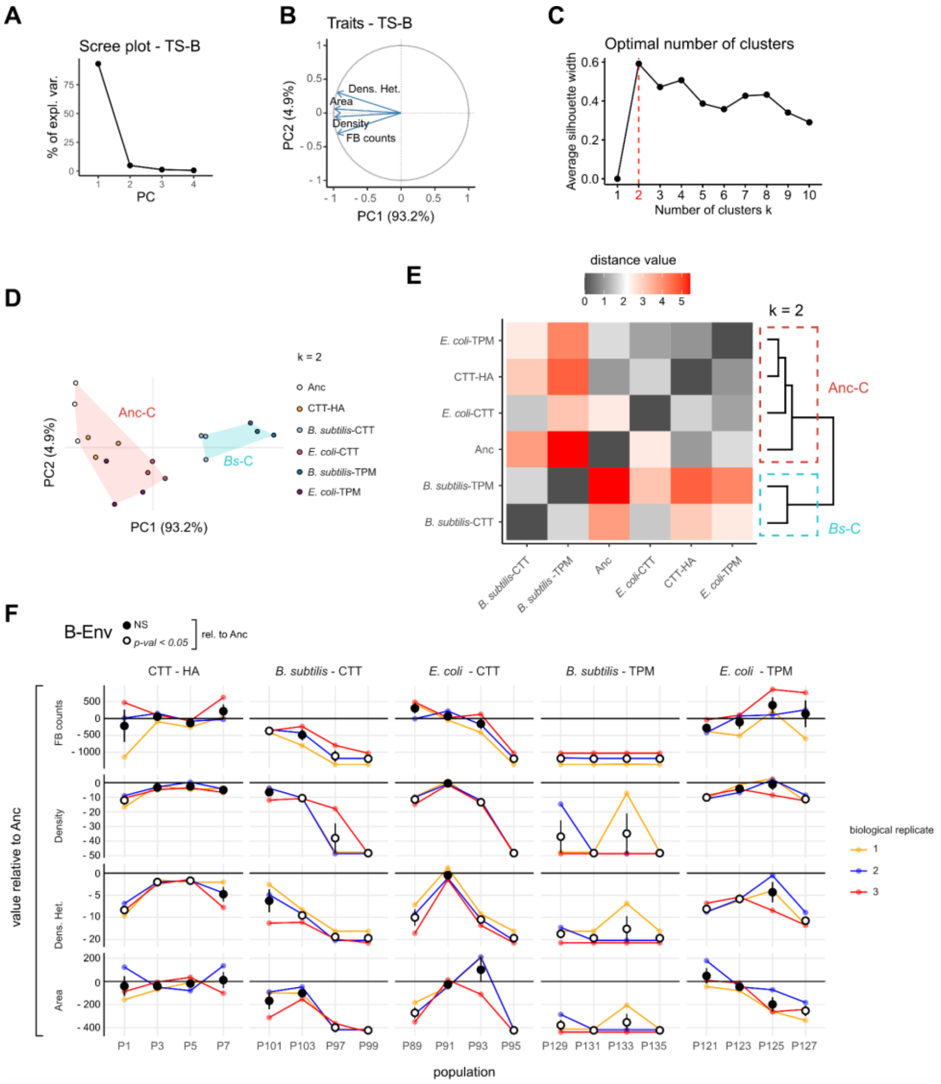


**Fig. S4. PCA, k-means, phenotype-dendrogram and population-level single traits analyses of TS-B.**

**A**, Scree plot indicating the percentage of the explained variance by each principal component obtained from the PCA run on TS-B and Anc together. **B**, Eigenvectors representing each of the four morphological traits used in the TS-B PCA. **C**, Optimal number of clusters (k = 2, highlighted in red on the x-axis) obtained from the average silhouette width (see Methods) of the TS-B PCA results. **D**, Subdivision of all individual replicates by the two optimal statistical clusters Anc-C (Ancestor-cluster, red-shaded region) and Bs-C (*B. subtilis*-cluster, blue-shaded region) obtained from the k-means cluster analysis (k = 2, see Methods). **E**, Heatmap of pairwise Euclidean distances measured between individual centroids mapped on the multivariate space (Fig. 1B). Individual tail colours ranging from 0 (grey) to 5 (red) indicate distance values. Dendrogram of hierarchical clustering (see Methods) across all pairwise Euclidean distances is reported on the right side. To facilitate the contrast between the two clustering methods, k-means clusters Anc-C (red-dashed box) and Bs-C (light-blue dashed box) (see **D**) were superimposed on the dendrogram. **F,** single developmental trait values relative to Anc (horizontal-black line set at zero) of all single evolved populations from TS-B (**B**) selective environments. Values were averaged across three independent biological replicates (small colored circles with connecting lines, *n* = 3). Average values (large circles) are reported with the associated SEM in all cases. Colored lines connect different population values measured within the same biological replicate. In both panels, white and black filled circles indicate average values with a significant or non-significant difference from the Anc, respectively. In **F**, significance was calculated in all cases with one-way ANOVA followed by two-tailed Tukey tests. *p*-values of all comparisons of evolved treatments with Anc, as well as all pairwise comparisons between evolved treatments are reported in Data Fig. S4.

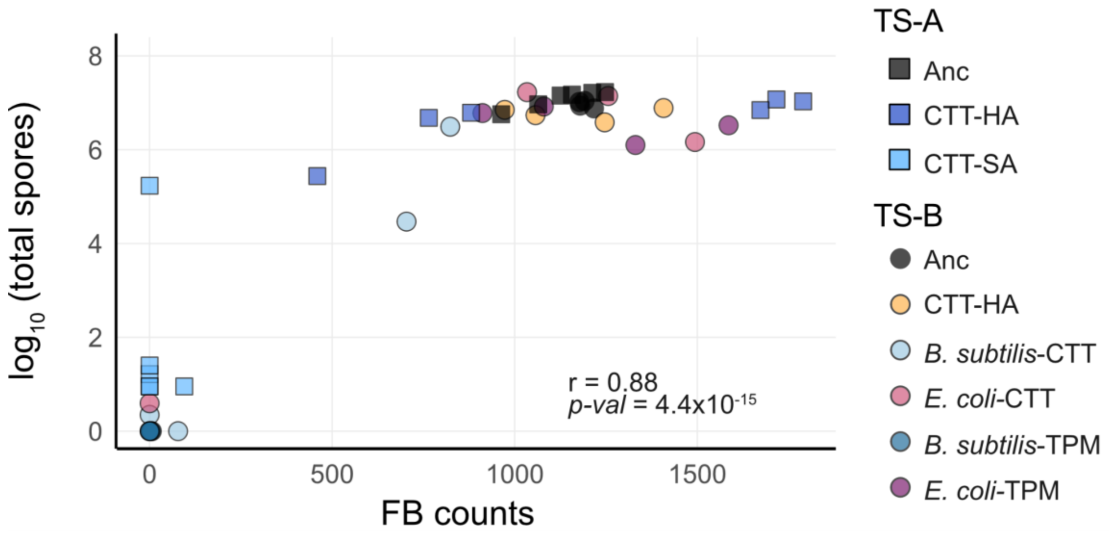


**Fig. S5. Fruiting body counts and spore counts correlate positively across all TS-A and TS-B populations and their ancestors.**

Scatterplot showing the relationship between average FB counts and the averaged log10-transformed spore counts across the three independent biological replicates for each individual TS-A and TS-B evolved population and the Anc sub-clones (n = 42). Pearson's correlation (r) and the associated *p*-value are shown inside the plot area. Data Fig. S5.

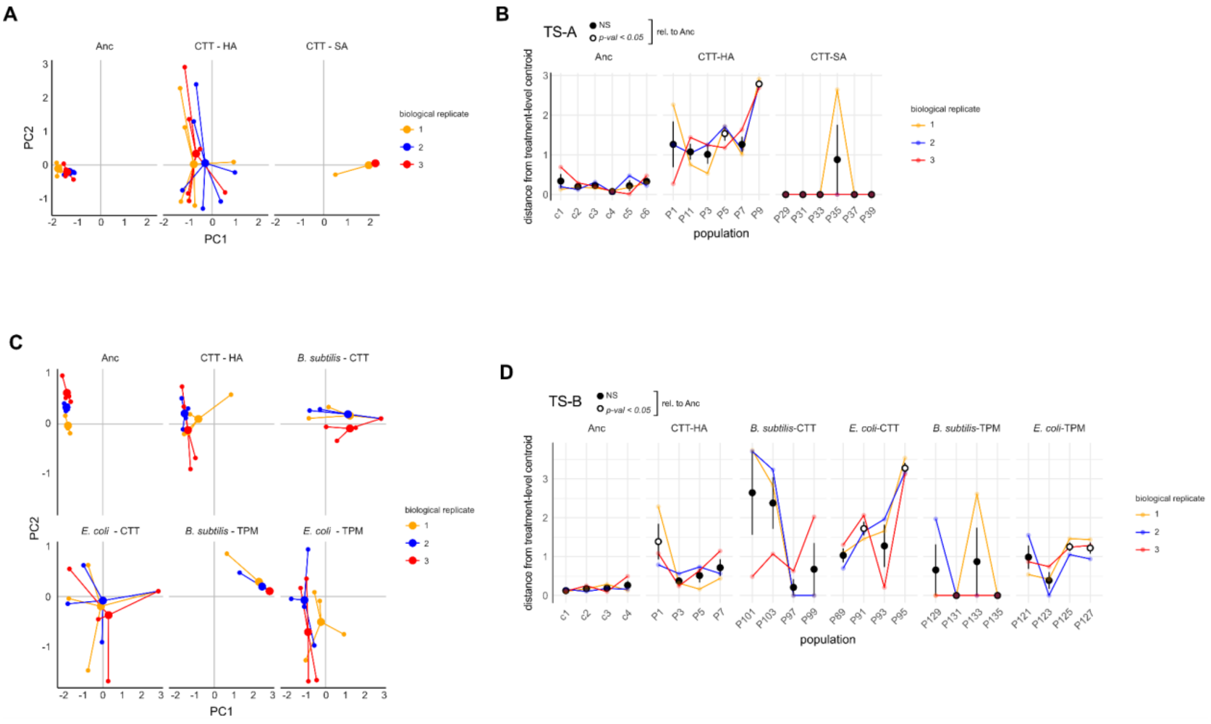


**Fig. S6. PCA-dispersion analyses among individual populations**

**A** and **C**, Morphospace showing the phenotypic distance (line lengths) of each population (small circles) from the averaged centroid (large circles) of all populations from the same treatment per replicate for TS-A (**A**) and TS-B (**C**) treatments. The three independent replicates are identified by color (orange, blue, and red). **B** and **D**, Quantitative plots (y axis) of the phenotypic distances reflected by line lengths in panels **A** and **C**. Average values per population across replicates are reported (large circles) with the associated SEM ($n$ = 3). Colored lines connect values of different populations measured within the same biological replicate (same color). For both panels **B** and **D**, white and black filled circles indicate average values with a significant or non-significant difference from the Anc average value, respectively (Significance was calculated in all cases with one-way ANOVA followed by two-tailed Tukey test. $p$-values of all comparisons of evolved treatments with Anc, as well as all pairwise comparisons between evolved treatments are reported in **Data Fig. S6**.

**Table S1. List of the MyxoEE-3 treatments relevant to this study (adapted from Table S2 in *Rendueles et al.*[74])**

| | | | **Examined populations** organized by ancestral GJV1 (Anc) sub-clone | | | | | |
|---|---|---|---|---|---|---|---|---|
| **Treatment** | **Description** | **MyxoEE-3 cycle** | GJV 1.1[a] | GJV 1.2 | GJV 1.3 | GJV 1.4 | GJV 1.5 | GJV 1.6 |
| ***Treatment Set A (TS-A)*** | | | | | | | | |
| CTT hard agar (CTT-HA) | 1% Casitone, 1.5% agar | 40 | P1 | P3 | P5 | P7 | P9 | P11 |
| CTT soft agar (CTT-SA) | 1% Casitone, 0.5% agar | 40 | P29 | P31 | P33 | P35 | P37 | P39 |
| ***Treatment Set B (TS-B)*** | | | | | | | | |
| CTT-HA | 1% Casitone, 1.5% agar | 18 | P1 | P3 | P5 | P7 | | |
| *E. coli*-CTT-HA here '*E. coli*-CTT' | *E. coli* grown on CTT HA | 18 | P89 | P91 | P93 | P95 | | |
| *B. subtilis*-CTT-HA here '*B. subtilis*-CTT' | *B. subtilis* grown on CTT HA | 18 | P97 | P99 | P101 | P103 | | |
| *E. coli*-TPM-HA here '*E. coli*-TPM' | *E. coli* overlaid on TPM HA | 18 | P121 | P123 | P125 | P127 | | |
| *B. subtilis*-TPM-HA here '*B. subtilis*-TPM' | *B. subtilis* overlaid on TPM HA | 18 | P129 | P131 | P133 | P135 | | |

[a] GJV1.1-GJV1.6 are labeled as c1-c6 in Fig. S6B,D.

**Table S2. Experimental differences between TS-B treatments during MyxoEE-3**

| | CTT-HA | *B. subtilis*-CTT | *E. coli*-CTT | *B. subtilis*-TPM |
|---|---|---|---|---|
| ***B. subtilis*-CTT** | prey presence (*Bs*) | - | - | - |
| ***E. coli*-CTT** | prey presence (*Ec*) | prey identity | - | - |
| ***B. subtilis*-TPM** | prey presence<br>casitone presence | casitone presence<br>prey-growth conditions[a] | prey identity<br>casitone presence<br>prey-growth conditions | - |
| ***E. coli*-TPM** | prey presence<br>casitone presence | prey identity<br>casitone presence<br>prey-growth conditions | casitone presence<br>prey-growth conditions | prey identity |

[a] *prey-growth conditions* refer to conditions for the period of overnight prey growth to stationary phase immediately prior to inoculation of MyxoEE-3 agar plates with *M. xanthus.* Prey growth during this period occurred either on the MyxoEE-3 agar plates themselves in the case of the *B. subtilis*-CTT and *E. coli*-CTT treatments or in CTT liquid (with an equivalent amount of casitone available as in the prey-CTT plates) prior to spreading on TPM starvation agar in the *B. subtilis*-TPM and *E. coli*-TPM treatments (see Methods).

**Table S3: Loci with mutations in clones from two or more of the TS-A populations examined here.** Populations shown in red exhibited a significant decrease in FB counts than the ancestor.

| Old locus ID | New locus ID | annotated function | CTT-HA | CTT-SA |
|---|---|---|---|---|
| *MXAN_6012* | *MXAN_RS29170* | response regulator | P1, P3, P5, P7 | P31, P33, P35, P37 |
| *frzF (MXAN_4138)* | *MXAN_RS20090* | protein methyltransferase FrzF | P1, P3, P5, P9, P11 | P29, P31, P33, P35, P37, P39 |
| *MXAN_5852* | *MXAN_RS28370* | sensory box histidine kinase | - | P31, P33, P35, P37 |
| *lonD (MXAN_2017)* | *MXAN_RS09780* | ATP-dependent protease LonD | - | P29, P33, P35, P37 |
| *MXAN_7216* | *MXAN_RS34935* | ICE-like protease (caspase) p20 domain protein | P3, P7, P9, P11 | - |
| *hsfB (MXAN_5365)* | *MXAN_RS26035* | response regulator/sensor histidine kinase HsfB | P1, P5, P9 | - |
| *MXAN_5032* | *MXAN_RS24445* | efflux transporter, HAE1 family, inner membrane component | P5 | P35, P37 |
| *MXAN_5030* | *MXAN_RS24435* | efflux transporter, HAE1 family, outer membrane efflux protein | - | P31, P33 |
| *MXAN_0289* | *MXAN_RS01415* | putative membrane protein | P3, P5 | - |
